# Intermolecular interactions play a role in the distribution and transport of charged contrast agents in a cartilage model

**DOI:** 10.1101/591255

**Authors:** Jenny Algotsson, Peter Jönsson, Jan Forsman, Daniel Topgaard, Olle Söderman

**Affiliations:** Division of Physical Chemistry, Lund University, Lund, Sweden; Division of Theoretical Chemistry, Lund University, Lund, Sweden

## Abstract

The transport and distribution of charged molecules in polyelectrolyte solutions are of both fundamental and practical importance. A practical example, which is the specific subject addressed in the present paper, is the transport and distribution of charged species into cartilage. The charged species could be a contrast agent or a drug molecule involved in diagnosis or treatment of the widespread degenerative disease osteoarthritis, which leads to degradation of articular cartilage. Associated scientific issues include the rate of transport and the equilibrium concentrations of the charged species in the cartilage and the synovial fluid. To address these questions, we present results from magnetic resonance micro-imaging experiments on a model system of articular cartilage. The experiments yield temporally and spatially resolved data on the transport of a negatively charged contrast agent (charge = −2), used in medical examinations of cartilage, into a polyelectrolyte solution, which is designed to capture the electrostatic interactions in cartilage. Also presented is a theoretical analysis of the transport where the relevant differential equations are solved using finite element techniques as well as treated with approximate analytical expressions. In the analysis, non-ideal effects are included in the treatment of the mobile species in the system. This is made possible by using results from previous Monte Carlo simulations. The results demonstrate the importance of taking non-idealities into account when data from measurements of transport of charged solutes in a system with fixed charges from biological polyelectrolytes are analyzed.

## Introduction

The function of articular cartilage is to provide low friction and wear resistance, as well as to distribute load in load-bearing joints, for instance, in hips and knees [1]. The optimal functioning of the joint depends on the structure, composition as well as the integrity of the extracellular matrix in cartilage, which is mainly composed of water, collagen and proteoglycans [1, 2]. The load resistance property of cartilage is largely dependent on the proteoglycan content, originating from the glycosaminoglycan (GAG) side chains of the proteoglycan [3]. The GAG is highly negatively charged on account of its carboxyl and sulfate groups that are ionized under physiological conditions. These negative charges are fixed within the extracellular matrix and thus the expression fixed charge density (FCD) of the cartilage is used. Please note that the value of FCD is negative.

Osteoarthritis is a degenerative disease that affects the function of cartilage. The disease is a common cause of disability and it is therefore of importance to understand its progression and to develop efficient treatments. In early stages of the disease, a loss of GAG occurs [4] and one method used to monitor the resulting decrease in the FCD is delayed gadolinium-enhanced magnetic resonance imaging of cartilage (dGEMRIC) [5]. This method utilizes magnetic resonance imaging (MRI) together with an intravenous injection of a contrast agent. The contrast agent commonly used is Gd(DTPA)^2–^ which is negatively charged and contains the paramagnetic ion Gd^3+^, which has a concentration dependent influence on the spin-lattice relaxation time, *T*_1_, of the ^1^H of the water [6]. The dGEMRIC method is based on the fact that the Gd(DTPA)^2–^ complex will, due to its negative charge, distribute in an inverse relation to the GAG content of the cartilage. This means that where there is a high concentration of GAG, indicating a healthy cartilage, there will be a low concentration of contrast agent, and vice versa. Therefore, a measure of the concentration of the Gd(DTPA)^2–^, and consequently a proxy for the GAG content of cartilage, is obtained by measuring *T*_1_ of the water. In passing, we note that the dGEMRIC method is used both *in vitro* and *in vivo* [5, 7–12]. The method has found most of its applications in *in vitro* studies and this will most likely continue to be the most important application.

dGEMRIC is dependent on the transport of Gd(DTPA)^2–^ from the site of the injection to the joint of interest, and, subsequently, on the diffusion controlled transport into cartilage, since cartilage lacks direct blood supply. It is therefore important to understand the physiochemical mechanisms that govern the transport and partitioning of Gd(DTPA)^2–^ in cartilage. The transport of the contrast agent will, for example, be affected by the chemical potential difference between the cartilage and the synovial fluid and, in this respect, the FCD of the cartilage plays an important role. Furthermore, the transport will be affected by the amount of collagen and glycosaminoglycan on account of steric effects [13–17]. This is an example of the general and important problem of diffusion of small molecules in a complex micro-heterogeneous environment, where crowding and excluded volume effects as well as electrostatic effects are important. The results presented here are therefore applicable to a wider class of related problems than just issues dealing with cartilage.

A previously developed magnetic resonance micro-imaging (*µ*MRI) setup for the investigation of features related to the dGEMRIC method in a model system of cartilage was used to investigate the questions raised above under controlled conditions [18]. The *µ*MRI setup gives spatially and temporally resolved information on the transport of Gd(DTPA)^2–^ in the model system, which is designed with the goal of investigating the effects of the electrostatic interactions in cartilage. To this end, the cartilage is represented by a polyelectrolyte solution and the synovial fluid is represented by a NaCl solution. A semipermeable membrane is used to separate the polyelectrolyte from the salt solution. The experimental results are compared with numerical solutions obtained using the finite element method (FEM) of the relevant equations describing the transport of ionic species in our system. Since previous studies have shown that ions in cartilage do not behave ideally [19–21], the influence of intermolecular interactions are included in the theoretical work presented in this paper. This is made possible by comparing the obtained data with previous Monte Carlo simulations on the model system [21]. Furthermore, the low concentration of Gd(DTPA)^2–^ makes it possible to describe the transport with an approximate analytic solution for a system without boundaries.

In summary, our research goal is to address the transport of charged molecules in complex micro-heterogeneous systems and to develop methods where intermolecular interactions are accounted for.

## Model system

The model system is outlined in Fig 1A-C. There is a polyelectrolyte solution at *z* < 0 and a salt solution at *z >* 0, separated by a semipermeable membrane at *z* = 0. The charged polyelectrolyte cannot pass through the membrane into the salt solution, which leads to a difference in electric potential on the two sides of the membrane, which in turn will be compensated for by a concentration difference of ions. The following scenario with respect to time, *t*, is investigated:

**Fig 1.**
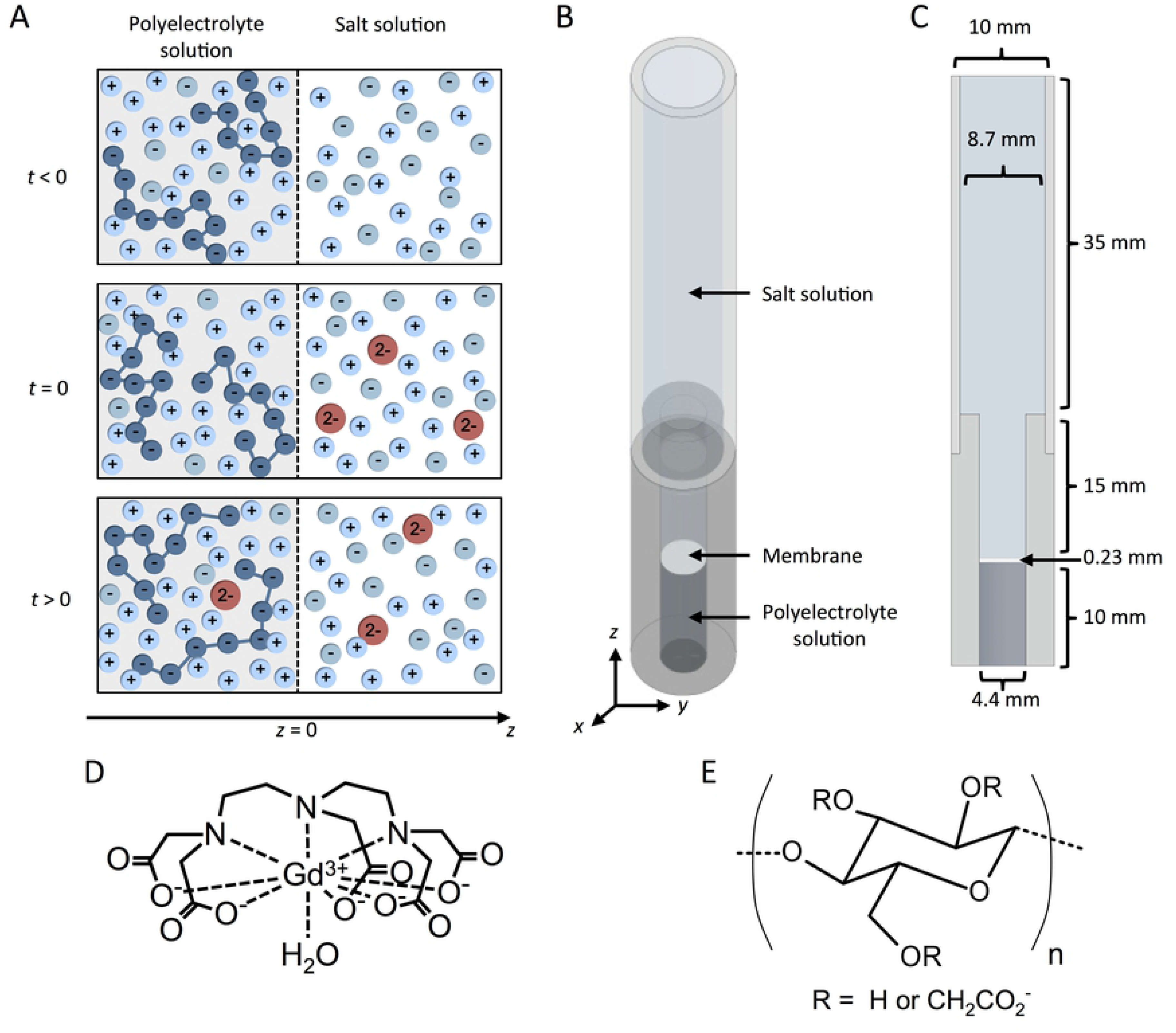
Presentation of the experimental and theoretical model systems. **(A)** The model system at different times. The polymer is negatively charged. At *t* < 0 there is thermodynamic equilibrium between the two solutions. At time *t* = 0, Gd(DTPA)^2–^ is added to the salt solution and is allowed to flow over the semipermeable membrane into the polyelectrolyte solution. In the system 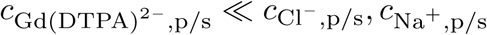, FCD (the subscripts p and s denote polyelectrolyte- and salt-solution, respectively). Moreover, because the polymer is negatively charged, the following will be true: 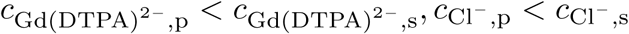 and 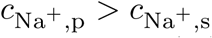. **(B)** Schematic drawing of the sample holder used in the *µ*MRI experiments. **(C)** Cross section of the sample holder in the x-y plane with indicated dimensions. **(D)** Chemical structure of Gd(DTPA)^2–^ [6]. **(E)** Chemical structure of carboxymethylcellulose (CMC).

1. At *t* < 0, there is equilibrium of Na^+^ and Cl^−^ ions between a salt solution and a solution containing a negatively charged polyelectrolyte.
2. At *t* = 0, a small amount of Gd(DTPA)^2–^ (see Fig 1D for the chemical structure of the complex) is added to the salt solution. The concentration of Gd(DTPA)^2–^ is much lower than the concentration of the other charged species in the system.
3. At *t >* 0, the Gd(DTPA)^2–^ will diffuse into the polyelectrolyte solution.

The concentration difference of Gd(DTPA)^2–^ between the solutions at *t >* 0 will lead to transport of the Gd(DTPA)^2–^ into the polyelectrolyte solution. Initially, the difference is large and leads to fast transport by mutual diffusion from the salt solution into the polyelectrolyte solution (a flow of Na^+^ and Cl^−^ ions will also occur to maintain the electro-neutrality in the system, which is minor compared to the total concentration of Na^+^ and Cl^−^ ions). As equilibrium is approached, the rate of transport decreases. Since the transport is only driven by diffusion, it will take a significant length of time to reach equilibrium. Recall that the root mean square displacement is proportional to the square root of time for diffusive motion.

## Theoretical background

The basic equations used in the interpretation of the data are presented in this section. The full derivation of the relations used in the Results and Discussion section is presented in the accompanying Supporting Information (SI).

The diffusive flux of an ionic species *i, J*_*i*_, occuring during step 3 above, can be described by the generalised Fick’s first law (in one dimension):

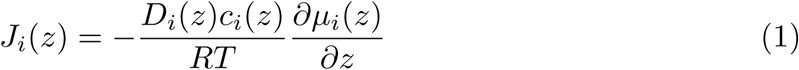

where *D*_*i*_ is the diffusion coefficient and *c*_*i*_ the concentration of ion species *i, R* is the gas constant and *T* the temperature. Furthermore, *µ*_*i*_ is the (electro)chemical potential of ion species *i*, which can, in turn, be expressed by the following equation:

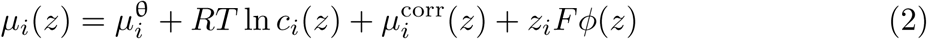

where 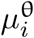 is the standard chemical potential, 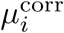 a correction factor for the chemical potential (resulting from intermolecular interactions), and *z*_*i*_ the charge of ion species *i*. *F* is the Faraday constant and *ϕ* the electric potential. If the expression for *µ*_*i*_ in Eq 2 is inserted into Eq 1, we obtain:

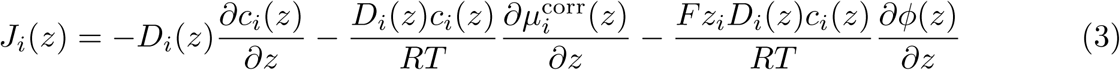

Moreover, the change of ion concentration with time, *t*, can be obtained from the equation of continuity:

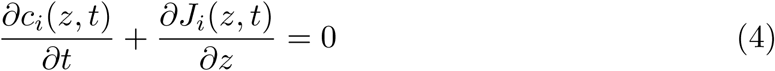

The electric potential, *ϕ*, is related to the charge distribution through Poisson’s equation:

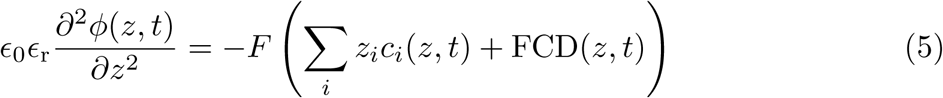

where *ϵ*_0_ is the permittivity of vacuum, *ϵ*_r_ is the dielectric constant and FCD is the fixed charge distribution, where FCD = 0 in the salt solution (*z >* 0) and non-zero in the polyelectrolyte solution (*z* < 0). In the present system, the concentration of Gd(DTPA)^2–^ is 2 to 3 orders of magnitude lower than the concentration of Na^+^ and Cl^−^ and, as a consequence, Eq 5 is reduced to:

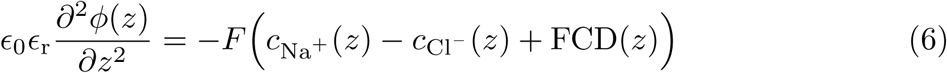

and the electric potential will thus be approximately constant for *t >* 0 on account of the fact that the relative changes in concentrations of Na^+^ and Cl^−^ are small at these times.

To be able to solve Eqs 3 and 4, 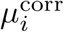 also needs to be known. It can be estimated by using results from previously made Monte Carlo simulations performed on a system developed to capture the electrostatic interactions in the cartilage/synovial fluid system at equilibrium [21]. In the simulations, the following holds for the two separate solutions, if the electric potential in the salt solution is set as the reference:

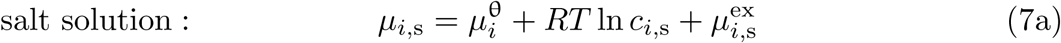

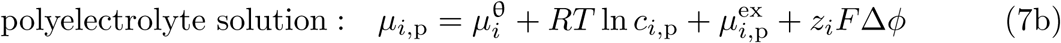

where 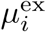 is the the excess chemical potential for ion *i* (related to the activity coefficient, *γ*_*i*_, according to: 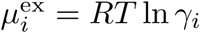) that is resulting from intermolecular interactions (i.e. long-ranged electrostatic interactions and repulsion because of steric overlap of particles), Δ*ϕ* is the difference in the electric potential between the two solutions, also known as Donnan potential, and subscripts s and p denote salt solution and polyelectrolyte solution, respectively. The simulations represent a situation of equilibrium, i.e. *µ*_*i*,s_ = *µ*_*i*,p_. The simulations also show that 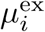 could approximately be seen as constant in the system over the steps 1, 2 and 3 above since, at low concentration of Gd(DTPA)^2–^, the intermolecular interactions between Gd(DTPA)^2–^ and Gd(DTPA)^2–^ ions can be ignored.

## Material and Methods

### Materials

Sodium carboxymethylcellulose (CMC) (average molecular weight = 90 kDa, 99.5% purity, Sigma Aldrich; molecular structure is given in Fig 1E) was used without further purification. The degree of substitution of the used batch of the polyelectrolyte has previously been determined to 0.9 [18]. Magnevist^®^ (0.5 mmol gadolinium/mL, Bayer Healthcare) was used as the Gd(DTPA)^2–^ source. The molecular structure of Gd(DTPA)^2–^ is given in Fig 1D. NaCl (99.5% purity, Merck) and NaOH (analytical grade) were used as received. All solutions were prepared with MilliQ purified H_2_O. Regenerated cellulose acetate membranes with a cutoff of 5 kDa were obtained from Millipore. This cutoff was previously shown not to affect the transfer rate of Gd(DTPA)^2–^ significantly and at the same time only allow for a negligible amount of CMC to pass through the membrane [18]. For dialysis of the polyelectrolyte solutions, Spectra/Por^®^ Biotech Cellulose Ester (CE) Dialysis membrane (dialysis cutoff of 100-500 Da) was used.

### Sample preparation

The pH of the salt solution (150 mM NaCl) was adjusted to between 8 and 9 by means of drop-wise addition of 1 M NaOH.

Magnevistt^®^ was diluted by the salt solution to a Gd(DTPA)^2–^ concentration of 0.36 mM. Before use, the pH was adjusted to the same value as in the salt solution.

When preparing the polyelectrolyte solutions, the CMC powder was dissolved in the salt solution (150 mM NaCl). Before use, the polyelectrolyte solutions (7 mL) were dialyzed for 24 h against 1.5 L 150 mM NaCl solution. A control experiment showed that this time was enough to reach ionic equilibrium between the polyelectrolyte solution and the salt solution (for more information, see the SI). The salt solution (the dialyzate) was gently stirred during the dialysis and the pH of the dialyzate was kept between 8 and 9 by periodically making drop-wise addition of 1 M NaOH. The carbon content in the polyelectrolyte solutions was determined by total organic carbon (TOC) analysis after the dialysis. The FCD was then calculated using as input the degree of substitution of the polyelectrolyte. To obtain further information with regard to the ionic content, the sodium content was determined in the solutions by inductively coupled plasma atomic emission spectroscopy (ICP-AES) analysis. Subsequently, the Cl^−^ concentration could be determined by applying electroneutrality conditions in the solutions. The densities of the solutions were determined by a precision density meter (DMA5000, Anton-Paar). The polyelectrolyte concentration of the polyelectrolyte solutions was estimated to 1.7, 2.2 and 2.5 wt%, respectively.

A small piece of the regenerated cellulose acetate membrane was punched out to fit into the sample holder (see Figs 1B and C). Before use, the membrane was, according to instructions from the manufacturer, put in 150 mM NaCl solution (with the glossy side up) to wash out glycerol and to wet the pores. The solution was changed three times during 1 h and thereafter, the membrane was left in the solution for at least 24 h before use.

### Experiments

The bottom section of the sample holder (constructed from polyether ether ketone (PEEK) plastic and a 10 mm NMR tube; see Figs 1B and C) was filled with *∼*160 *µ*L CMC solution (added by weight). Thereafter, the membrane was carefully placed on top of the polyelectrolyte solution with the glossy side down. Subsequently, the top part of the sample holder (including the NMR tube) was assembled with the bottom part and 205 *µ*L of the 150 mM NaCl solution was added on top of the membrane.

2.1 mL of the 0.36 mM Gd(DTPA)^2^ solution was added to the top compartment about 30 minutes after the addition of the salt solution (see above). The first *T*_1_ measurement (for description of *T*_1_ measurements, see below) was started within a few minutes after the addition of Gd(DTPA)^2–^. Due to rapid initial changes of the Gd(DTPA)^2–^ concentration, the initial *T*_1_ measurement (for each sample) gave unphysical concentration profiles in the salt solution and these data points were not used in the subsequent analysis. The sample holders were kept in the spectrometer for the initial 22 h and *T*_1_ measurements were performed continuously. For the following data points, the sample holders were removed and 3 measurements at different positions were collected (one showing the whole polyelectrolyte solution, one showing the polyelectrolyte and salt solutions at the same time and one showing the salt solution close to the membrane). The sample holder was inserted into the spectrometer for each individual measurement. This measurement procedure was carried out once a day. Between the measurements, the sample holders were stored at 298 K, protected from light. The samples were not stirred during the experiments and, in order to minimize agitation, the sample holders were carefully handled between measurements. Three different concentrations of CMC were studied and for each concentration, two independent experiments were performed.

### *µ*MRI measurements

Spatially resolved *T*_1_ measurements were recorded at 298 K on an 11.7 T Bruker Avance II 500 spectrometer equipped with a Bruker MIC-5 microimaging probe having a maximum gradient strength of 3 T m^−1^ and a 10 mm saddle-coil radio-frequency insert. The measurements were performed using a pulse sequence containing an inversion recovery block followed by a rapid acquisition with relaxation enhancement (RARE) block [22] for the read out of the images. The recovery time delay, *τ*, was incremented logarithmically in 32 steps from 0.0017 s to 15 s, while the images were taken up with an 11.1 *×* 4.8 mm field-of-view (*z × y*), 128 *×* 48 acquisition matrix size and a 0.2 mm slice thickness (*x*). This resulted in a measurement time of about 10 minutes per image.

### Data processing

The images were reconstructed to a 128 *×* 48 matrix size, which gives a spatial resolution of 87 *×* 100 *µ*m, and subjected to 0.2 mm Gaussian smoothing. For each voxel, the *T*_1_ values were obtained by fitting the image intensity, *I*, to following equation:

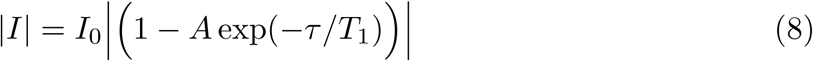

where *I*_0_ is the maximum image intensity, *τ* is the recovery time delay, and *A* is a constant, which is less than 2 if the pulse repetition time is less than around 5 times *T*_1_ or the inversion pulse is imperfect. Errors representing one standard deviation were obtained by performing Monte Carlo error estimations as described in [23].

The obtained *T*_1_ values were then converted to the concentration of Gd(DTPA)^2–^, 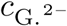, using the expression:

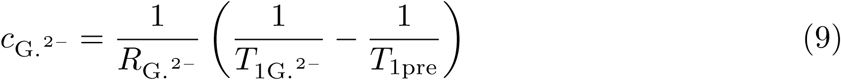

where 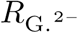 is the relaxivity of Gd(DTPA) ^2–^,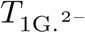 and *T*_1pre_ are the values of *T*_1_ with and without Gd(DTPA) ^2–^, respectively. 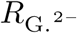 was determined in all used solutions (the salt solution and the three different polyelectrolyte solutions with different concentrations of CMC) by measuring *T*_1_ for different concentrations of Gd(DTPA)^2–^. The 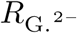 value in the salt solution was 3.84 *±* 0.03 (mM s)^−1^. The obtained values for the polyelectrolyte solutions are collected in Table 1. Errors in 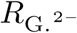 are estimated from the uncertainties in the obtained *T*_1_ values.

**Table 1.** Parameter values. Determined values for the different sets of experiments.

| FCD <sup>a</sup><br>(mM) | $C_{\text{NaCl},s}$ <sup>a</sup><br>(mM) | $c_{\text{Na}^+,p}$ <sup>a</sup><br>(mM) | $c_{\text{Cl}^-,p}$ <sup>a</sup><br>(mM) | $R_{\text{G},2-}$ <sup>a</sup><br>(mM<sup>-1</sup> s<sup>-1</sup>) | $D_{\text{G},2-,p}$ <sup>b</sup><br>(mM<sup>2</sup>/s) | $\Delta\mu_{\text{Na}^+}^{\text{corr}}$ <sup>c</sup><br>(RT) | $\Delta\mu_{\text{Cl}^-}^{\text{corr}}$ <sup>d</sup><br>(RT) | $\Delta\mu_{\text{G},2-}^{\text{corr}}$ <sup>e</sup><br>(RT) |
| --- | --- | --- | --- | --- | --- | --- | --- | --- |
| -73 | 150 | 202 | 129 | $3.92 \pm 0.02$ | $4.1 \cdot 10^{-10}$ | -0.0958 | -0.0512 | -0.2050 |
| -92 | 150 | 216 | 124 | $3.94 \pm 0.04$ | $3.9 \cdot 10^{-10}$ | -0.1105 | -0.0635 | -0.2600 |
| -108 | 150 | 223 | 115 | $3.96 \pm 0.02$ | $3.7 \cdot 10^{-10}$ | -0.1211 | -0.0099 | -0.2500 |

<sup>a</sup>For determination, see Material and Methods. <sup>b</sup>Determined by fitting of the experimental results with Eq S21 in the interval 2.5 - 6h. <sup>c</sup>From Monte Carlo simulations. <sup>d</sup>From Eq S24. <sup>e</sup>From FEM simulations, see text for details.

The three different images that were taken at the same time point (see above) were visually merged to obtain a larger field-of-view of the system.

Assuming that there is no convection in the polyelectrolyte solution, the diffusion coefficients of Gd(DTPA)^2–^ in the polyelectrolyte solutions were obtained from fitting the relevant equations to the concentration profiles in the polyelectrolyte solution (see the SI for explanation of the procedure). The data points used in the fit were obtained in a time interval where the influence of the membrane is low and the Gd(DTPA)^2–^ has not yet reached the bottom of the polyelectrolyte compartment, i.e. 2.5-6 h.

All data processing was performed with in-house written Matlab^®^ code.

### Numerical Methods

Numerical calculations were carried out using COMSOL Multiphysics^®^ 4.4, a program that solves partial differential equations using the finite element method (FEM). The FEM simulations were carried out for a one-dimensional geometry with the same dimensions as in Fig 1C, except that the salt solution was truncated at *z* = *d*_salt_ to account for thermally driven convection in the salt solution. At this distance, the concentration in the salt solution in the FEM simulations was set equal to the experimental value at the same position. The distance *d*_salt_ was varied between 1 to 5 mm for the different experiments, depending on whether the contribution from convection was high or low, respectively. This was determined by comparing the experimental concentration profiles in the salt solution with those obtained from the FEM simulations without convection and choosing *d*_salt_ at a value where the experimental and the calculated curves in the salt solution are equal. Moreover, we assume homogenous properties in each solution, such that values of the diffusion coefficients are constant in each solution and the chemical potential correction factors are, except close to the interface, also constant in each solution.

Data from all experiments (two sets of data for each polyelectrolyte concentration) were used in the simulations. First, the time-independent Eq 4 and Poisson’s equation (Eq 6) were solved for the Na^+^ and Cl^−^ ions as well as for the electric potential. In order to avoid a singularity in the change in chemical potential at *z* = 0, we introduced a smooth change in 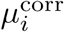 around *z* = 0 according to:

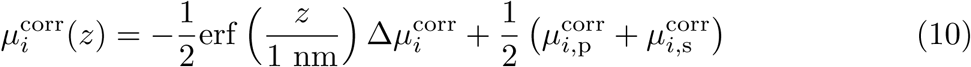

where 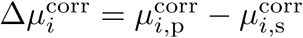 (in what follows, the second indices p, and s denote polyelectrolyte and salt solutions, respectively). Choosing a different value than 1 nm for the range over which the chemical potential change has negligible effect on the macroscopic difference in electric potential between the polyelectrolyte and salt solution, Δ*ϕ*, and thus according to Eq S2 does not affect the obtained value for the macroscopic step in concentration of Gd(DTPA)^2–^ at *z* = 0. The obtained values of 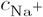, 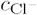 and *ϕ* were subsequently used to calculate the distribution of Gd(DTPA)^2–^ as a function of time by solving Eqs. 3 and 4 for Gd(DTPA)^2–^. The FEM simulations were run for each experiment between 80 h and 260 h.

The boundary conditions used are summarized in Table 2. The initial concentrations were 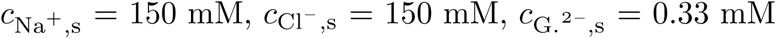 and 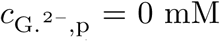. The temperature was 298.15 K, *ϵ*_r_ = 78, and the diffusion coefficients for Na^+^ and Cl^−^ were assigned values of 1.33 · 10^−9^ m^2^/s and 2.03 · 10^−9^ m^2^/s, respectively [24]. It should be noted that the obtained results are independent of the chosen diffusion coefficients, since we are assuming a situation of steady state for the Na^+^ and Cl^−^ ions. The value for the diffusion coefficient for Gd(DTPA)^2–^ used in the salt solution was obtained from a study of molecules similar to Gd(DTPA)^2–^ and was taken to be 4.5 · 10^−10^ m^2^/s [25]. The diffusion coefficient for Gd(DTPA)^2–^ in the membrane was set to 1.5 · 10^−10^ m^2^/s in all FEM simulations, chosen to fit with the experimental data at early measurement times. As noted above, the diffusion coefficients in the polyelectrolyte solution were obtained from analysis of the experimental data (see also below and in the SI). Values for 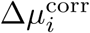 were determined from the equilibrium concentrations together with equations describing equilibrium concentrations in the framework of a Donnan equilibrium. However, values for 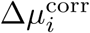 cannot be determined independently for Na^+^ and Cl^−^ as there are two independent relations and three unknowns (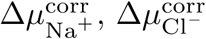 and 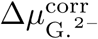 see Eq S24). To arrive at individual values of 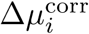 needed in the analysis of Gd(DTPA)^2–^, we start by setting the 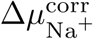 to values of 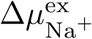 obtained from Monte Carlo simulations [21]. The 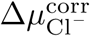 values are subsequently obtained from the equilibrium concentrations of Na^+^ and Cl^−^ (see Table 1) and Eq S24. In principle, values for 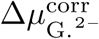 can then be determined from Eq S24c. However, the equilibrium concentrations of Gd(DTPA)^2–^ are less accurately known, on account of the long time required to reach equilibrium. To improve the accuracy in the obtained value, we have made an initial estimate from Eq S24c and then allowed the equilibrium value of Gd(DTPA)^2–^ to vary in the FEM simulations so that the simulated concentration profiles best fit the experimentally determined ones. The used values are collected in Table 1.

**Table 2.**
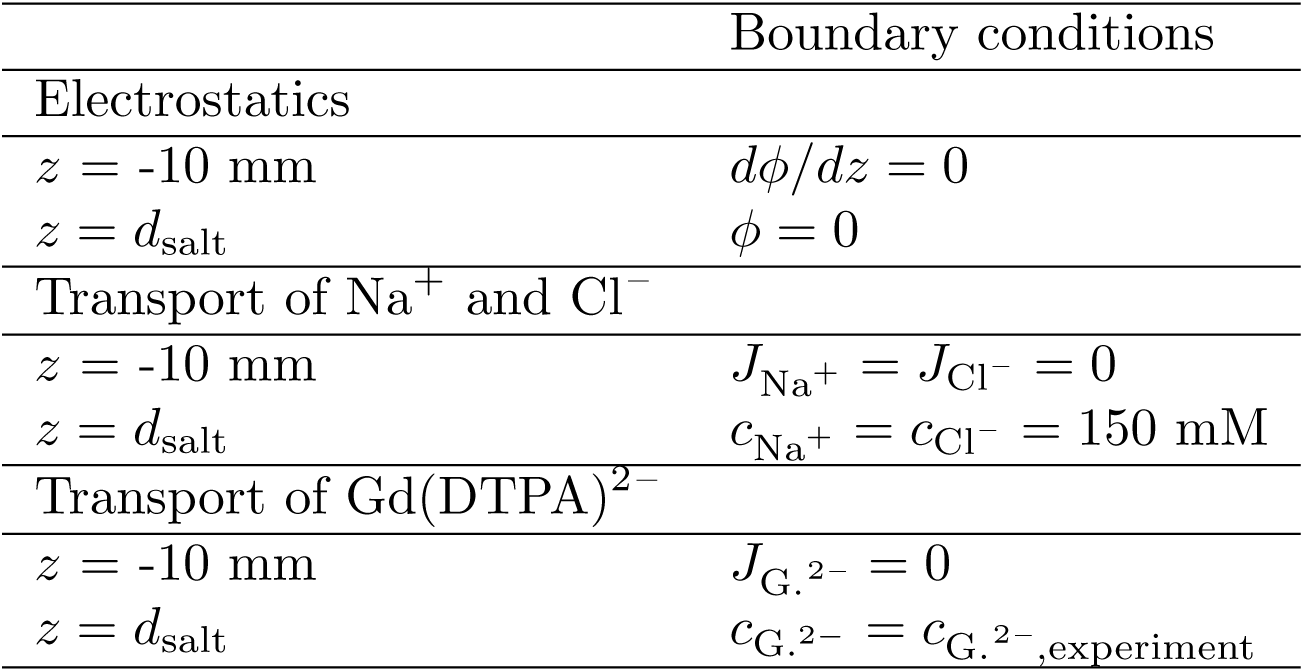
Boundary conditions. Boundary conditions used in the finite element method (FEM) simulations.

To investigate the validity of the approximate equations that are derived in the Results and Discussion section, FEM simulations were also carried out with same dimensions as in Fig 1C (not truncated at a distance *d*_salt_ as described above) and values taken from Monte Carlo simulations (ref. 21 and unpublished results, see the SI).

## Results and Discussion

The outline of this section is as follows. We start by presenting and discussing the experimental data obtained, and then proceed to numerically solve the relevant equations describing the data using FEM. Subsequently, we develop approximate solutions that can be used to illustrate how different parameters affect the distribution of ions in the system. Using examples, we demonstrate under which conditions the approximate relations are valid. Finally, we use the approximate relations to determine the diffusion coefficients of the contrast agent in the polyelectrolyte solution. In the interest of keeping the paper short, we delegate derivations to the SI, to which we refer when needed.

### Experimental data

The main experimental results of this study are concentration profiles for Gd(DTPA)^2–^ as a function of time for three different values of FCD. An example is given in Fig 2, which shows the experimentally determined concentrations of Gd(DTPA)^2–^ in a system with the polyelectrolyte solution having a FCD of −108 mM, at different times after addition of Gd(DTPA)^2–^ to the salt solution (results for the duplicate experiment with FCD = −108 mM and the other values of FCD are shown in the SI). After the addition, Gd(DTPA)^2–^ diffuses from the salt solution to the polyelectrolyte solution. This is a slow process as can be observed in Fig 2. The times needed for the average concentration in the polyelectrolyte solution to reach 50% and 90% of the equilibrium concentration in the 10 mm long polyelectrolyte solution are 40 h and 230 h, respectively. However, there is, as expected, a rapid development of a discontinuity in the concentration at *z* = 0, although it appears to be gradual in Fig 2 due to the finite resolution of the *µ*MRI method and artifacts due to the presence of the membrane (e.g. uncertain values of relaxivities and *T*_1pre_). The FEM simulations make it possible to study the discontinuity closer (see below).

**Fig 2.**
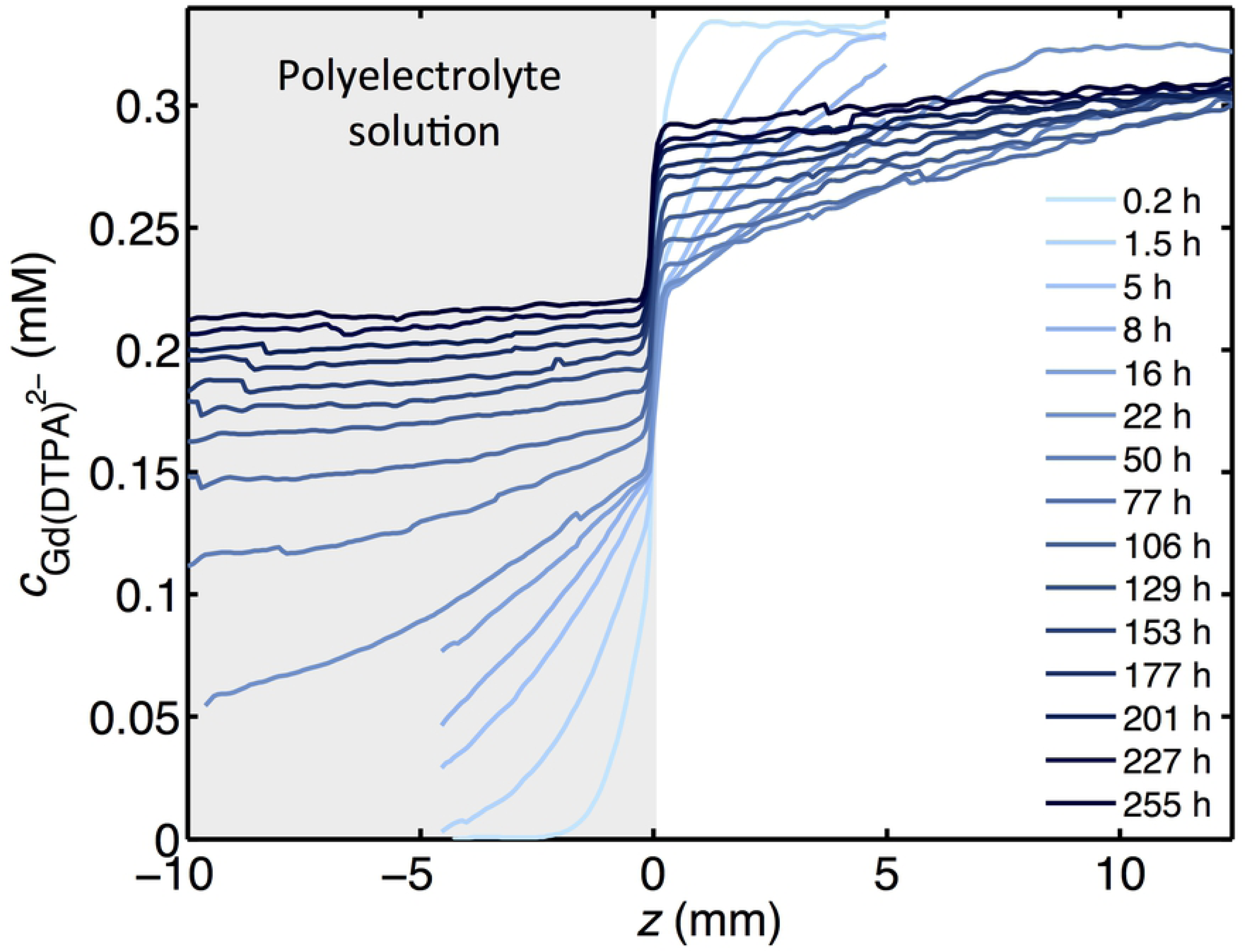
Experimental concentration profiles. Concentration profiles of Gd(DTPA)^2–^ at different times in the salt solution and the polyelectrolyte solution with FCD = −108 mM (*z* < 0). Gd(DTPA)^2–^ was injected in the salt solution at *t* = 0 and the data was acquired using *µ*MRI. The (small) decrease of the concentration of Gd(DTPA)^2–^ at longer times on the right hand side is due to the finite volume of the salt solution.

In Fig 3A, we show the concentration of Gd(DTPA)^2–^ in the polyelectrolyte solution at a position 4 mm from the membrane as a function of time for a FCD of −108 mM. The concentration of Gd(DTPA)^2–^ will initially increase rapidly in the polyelectrolyte solution, but the magnitude of the increase will gradually decrease with time. The symbols present two sets of data recorded in repeated experiments with the same FCD for the polyelectrolyte solution. The difference in the curves is attributed to a different degree of temperature gradient driven convection in the salt solution at the beginning of the experiments. The contribution from convection can be estimated by analyzing the concentration profiles in the salt solution (see Fig 2A and the SI). Furthermore, convection evens out concentration gradients in the salt solution and therefore, the time to reach equilibrium becomes shorter. On account of the geometry (small radius and height) and the position of the sample holder in the spectrometer (where temperature gradients are small), and the fact that the polyelectrolyte is rather viscous, the transport in the polyelectrolyte solution is well described by pure diffusion.

**Fig 3.**
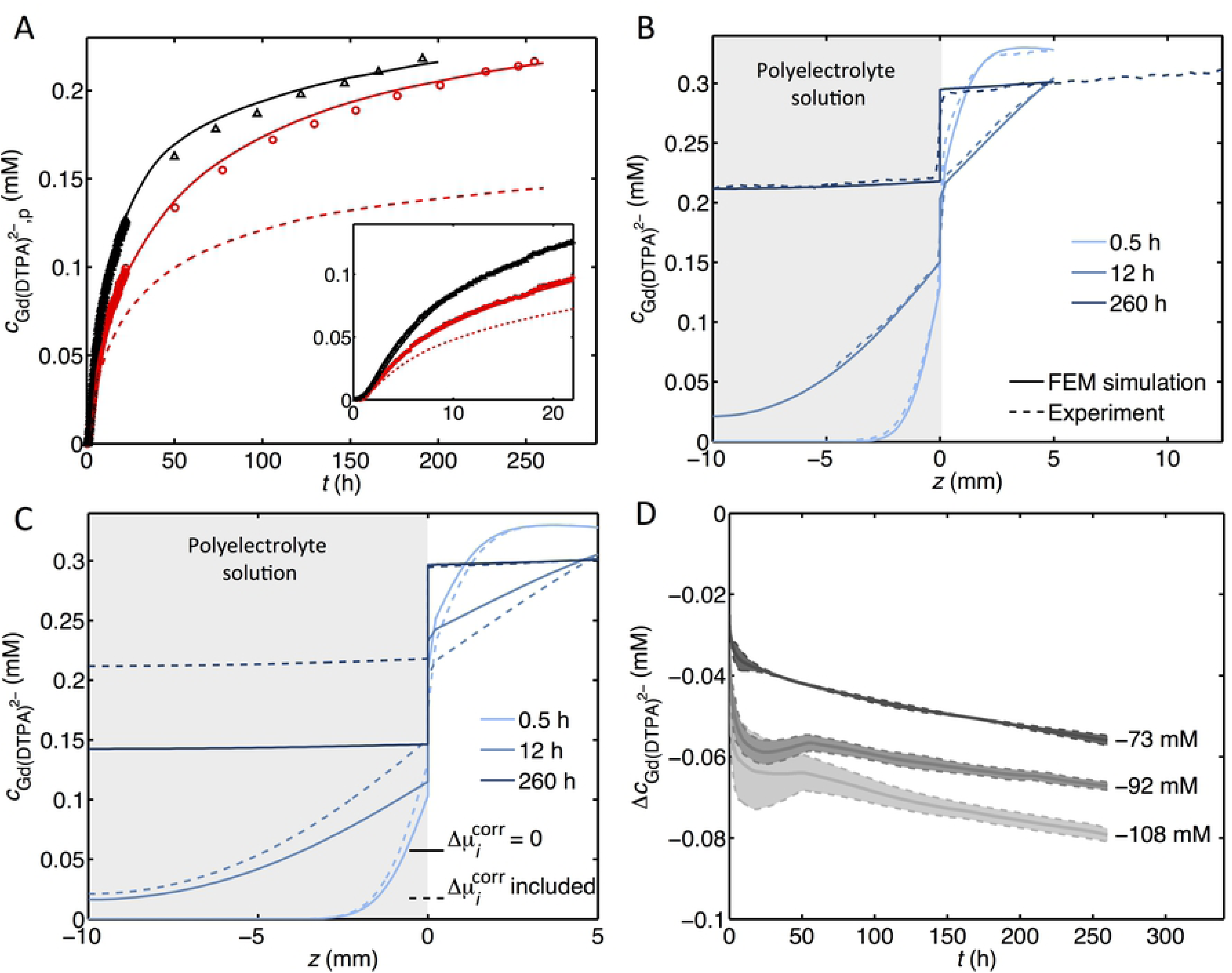
Compiled results from experiment and FEM simulations. **(A)** The concentration of Gd(DTPA)^2–^ in the polyelectrolyte solution at a position 4 mm from the membrane for two samples with FCD = −108 mM: one with small effects due to convection (circles) and one with noticeable convection at the beginning of the experiment (triangles), as a function of time. The solid lines are the corresponding values from FEM simulations with fitted values of 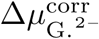 and the dashed line represents the corresponding results from a simulation with 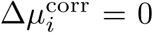 for all ions. **(B)** Experimental (dashed) and simulated (solid) values for the concentration of Gd(DTPA)^2–^ at different positions along the container, *z*, for a polyelectrolyte solution with FCD = −108 mM. **(C)** Simulated values for the concentration of Gd(DTPA)^2–^ at different positions along the container, *z*, for a polyelectrolyte solution with FCD = −108 mM. The dashed line shows the result from a FEM simulation with fitted value of 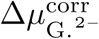 and the solid line represents the corresponding results from a simulation with 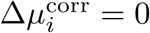 for all ions. **(D)** The step in concentration at *z* = 0, 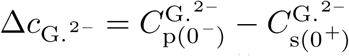 (see also Fig 4), at different times for three polyelectrolyte solutions with different FCD. The dashed lines correspond to the individual experiments and the solid lines represent the corresponding mean value of the duplicate experiments. The data are taken from FEM simulations.

In the following sections, we will use relevant theoretical expressions in combination with FEM simulations to investigate and rationalize the behavior of the Gd(DTPA)^2–^ with regard to its distribution and transport. The analysis yields physical properties relevant for the system. We note that the approach can be used to predict and understand the transport of ions in general in related systems.

### Finite element method simulations

By solving Eq 3, 4 and 6 numerically, using FEM techniques (as described above in Material and Methods), we are able to predict the experimental results in Fig 2. Using as input the data in Table 1 (and values for the diffusion coefficients for Na^+^ and Cl^−^ given above) and the boundary conditions in Table 2, the results are presented as solid lines in Figs 3A and 3B (for additional comparison between experiments and FEM simulations, see the SI). As can be seen, there is a good agreement with the experimental results at all times, and the effects due to convection in the salt solutions are well accounted for. We stress that all relevant parameters have been determined in independent experiments, except for the values of the electric potential and the chemical potential correction factor for the mobile ions, which are in good agreement with the results from independent Monte Carlo simulations (ref. 21 and unpublished results, see SI).

One of the key motivations for carrying out this study was to investigate to what extent effects due to non-ideal conditions are important in the transport and equilibrium concentrations of Gd(DTPA)^2–^ in cartilage and models of cartilage. This is one important issue in the dGEMRIC method. To this end, we present in Figs 3A and 3C results of FEM simulations where the difference in the chemical potential correction factors between the two solutions of all species are set to zero. It is clear that the predicted equilibrium concentrations deviate considerably, in agreement with earlier work [5, 18, 21].

The FEM simulations enable us to study the discontinuity in the concentration at *z* = 0. In Fig 3D, the step in the Gd(DTPA)^2–^ concentration 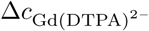 (see also Fig 4), is plotted against time. There is a rapid development of a discontinuity in the concentration at *z* = 0 and the magnitude of the step increases with time. The double experiments for each value of FCD, show that the magnitude of the step size is reproducible. The deviations are due to different degrees of convection in the salt solution. While the steps in concentration vary with time, the ratios between the concentrations of Gd(DTPA)^2–^ in the polyelectrolyte solution and salt solution at *z* = 0 are independent of time, with values of 0.82, 0.78 and 0.74 at FCD values of −73 mM, −92 mM and −108 mM, respectively.

**Fig 4.**
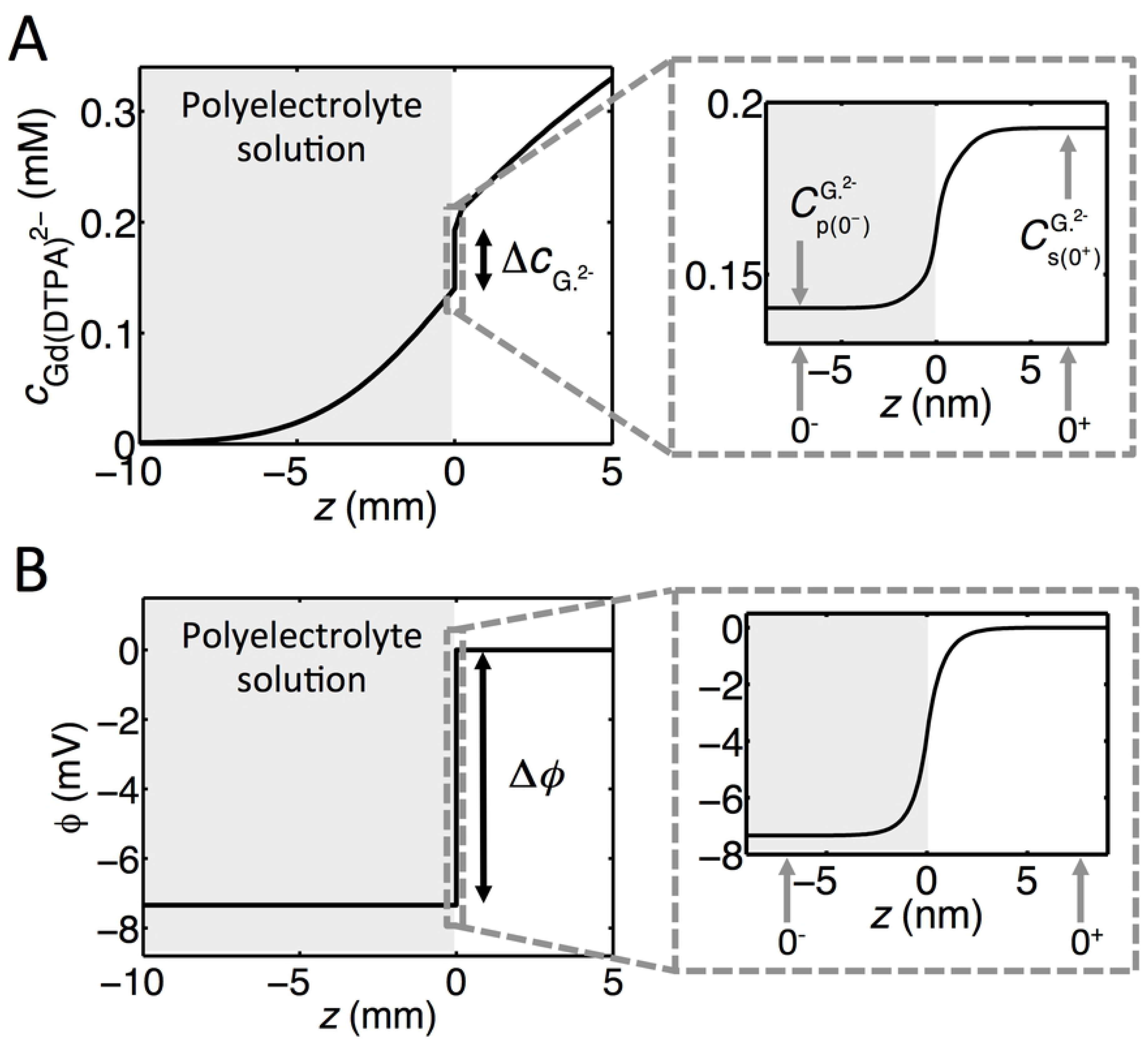
Concentration and potential profiles from FEM simulations. **(A)** A concentration profile for Gd(DTPA)^2–^ 5h after addition of Gd(DTPA)^2–^ in the salt solution. **(B)** The corresponding potential profile. The profiles are taken from FEM simulations with a polyelectrolyte solution with a FCD of −108 mM. The inserts to the right show the profiles around *z* = 0.

### Approximate relations for concentration profiles

Motivated by the need to illustrate the influence of the relevant parameters and to carry out rapid and straightforward calculations of the diffusion and concentration profiles, we here develop approximate expressions describing the transport and concentration profiles of Gd(DTPA)^2–^ in our model system. Our point of departure is to consider an interval 0 ^−^ ≤ *z* ≤ 0^+^ (see Fig 4), the size of which is such that we can assume that quasi steady-state conditions apply. This is justified by the fact that the gradient in the concentration at *z* = 0 adjusts much more rapidly than the variation in concentration over macroscopic length scales outside 0^−^ ≤ *z* ≤ 0^+^. In the current experiments, this interval is around 10 nm which is orders of magnitude smaller than the dimensions of the macroscopic container (see Fig 4). Solving Eq 3 with *J*_*i*_ = 0 gives (the reader is referred to the SI for details):

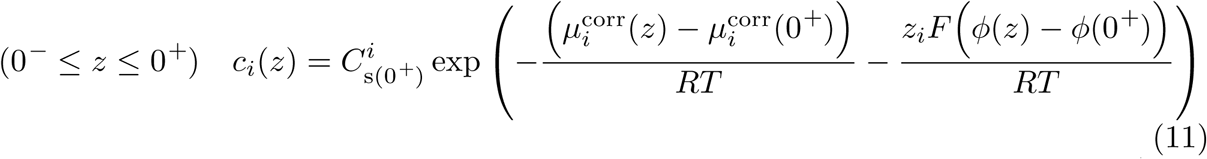

where 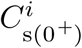 is the macroscopic concentration of ion *i* in the salt solution at *z* = 0^+^. To obtain approximate expressions for the step sizes in potential and concentration between the salt solution and the polyelectrolyte solution, Eq 11 is linearized (see Eq S4). Inserted into Eq 6, this gives the following, approximate expression for the electric potential drop between the polyelectrolyte solution and the salt solution:

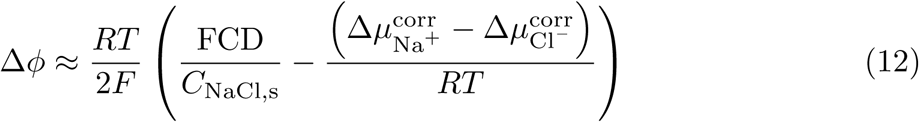

where Δ*ϕ* is the electric potential difference between the polyelectrolyte solution and the salt solutions, *C*_NaCl,s_ is the concentration of Na^+^ and Cl^−^ in the salt solution (the small difference in concentration due to the presence of Gd(DTPA)^2–^ ions is neglected) and 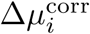 is defined above. We note that Ohshima and coworkers have analyzed a system that is similar to ours and present results which are in agreement to those presented here when excess chemical potentials are neglected. [26, 27] By combining Eqs 11 and 12 and linearizing the exponent, we obtain the following approximate expressions for the step in concentration, Δ*c*_*i*_, between the polyelectrolyte solution and the salt solution:

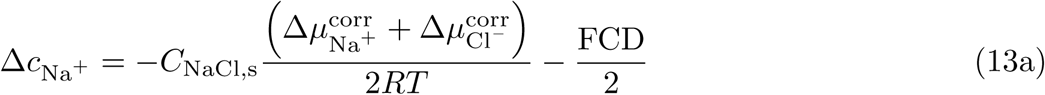

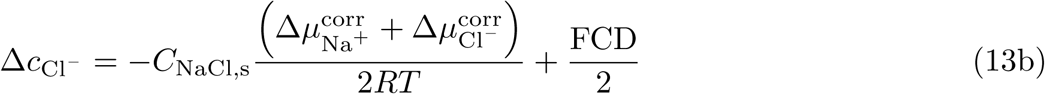

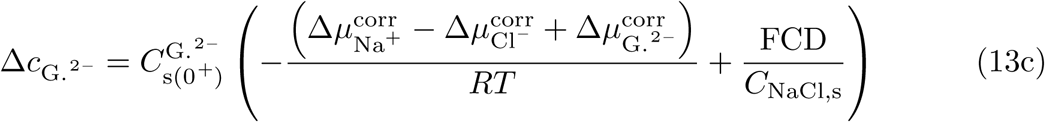

From Eq 13, it is clear that the difference in the chemical potential correction factor between the solutions changes the step in the concentration at the interface with the same amount for both Na^+^ and Cl^−^, and therefore, as noted above, the individual values of 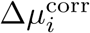 for the different ions cannot be determined from measurements of the concentration steps alone. The expressions above are only valid for small changes in the concentration (on account of the linearization), but they are useful in order to illustrate how the different parameters affect the distribution of ions in the system and qualitatively understand the behaviour of Fig 3D. Thus, the concentration of Gd(DTPA)^2–^ in the polyelectrolyte solution will decrease due to the negative value of the FCD of the polyelectrolyte solution. Furthermore, the difference in 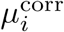 between the two solutions can either increase the concentration step, as for Na^+^, or decrease the concentration step, as for Cl^−^ and Gd(DTPA)^2–^ (cf Eqs 13 and Table 1). The step size will increase as Gd(DTPA)^2–^ reaches the bottom wall of the container, resulting in an increase in 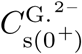, which is the reason for the increasing magnitude of the concentration step size observed in Fig 3D. Additionally, Eq 13 illustrates the fact that the assumption of 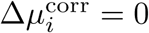 for all mobile ions, mentioned above, has a substantial impact on the step size at all times (cf. Fig 3C and Table 1).

We next carry out an approximate analysis of the transport of Gd(DTPA)^2–^. To this end, we solve Eq 13 on both sides of the salt/polyelectrolyte solution interface at *z* = 0

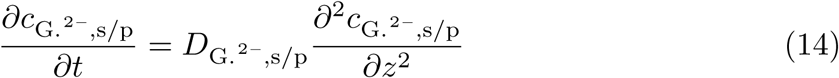

which is obtained by combining Eq 3 and Eq 4 and is valid outside the interval 0 ^−^ < *z* < 0^+^. The term containing the electric potential in Eq 3 can be neglected since the concentration of Na^+^ and Cl^−^ is much larger than the concentration of Gd(DTPA)^2–^ resulting in the electro-kinetic transport of the Gd(DTPA)^2–^ ions being small compared to the diffusional transport. The concentration difference between the salt solution and the polyelectrolyte solution is approximately given by Eq 13c, which together with the condition that the flux of ions on both sides of *z* = 0 should be equal, couples the solutions for the polyelectrolyte and salt solutions. The solution to these equations for an infinite system without convection and membrane is given by (see the SI for details):

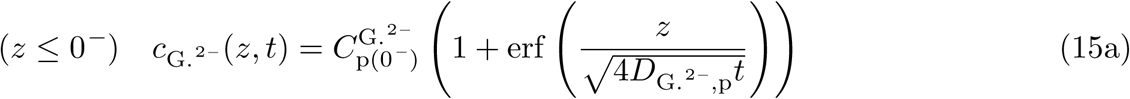

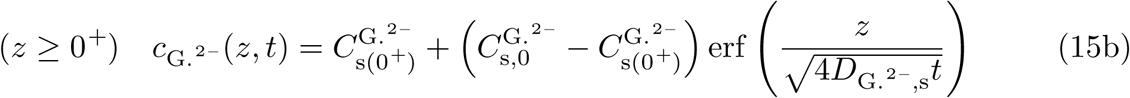

where 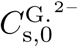 is the concentration of Gd(DTPA)^2–^ at *t* = 0 in the salt solution (*z* < 0) and 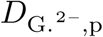 and 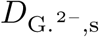 are the diffusion coefficients of Gd(DTPA)^2–^ in the polyelectrolyte solution and the salt solution, respectively. The concentrations 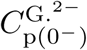 and 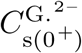 are related via Eq 13c, where 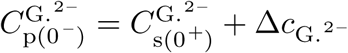. From the condition that the flux is equal at *z* = 0^−^ and at *z* = 0^+^, the following condition also applies:

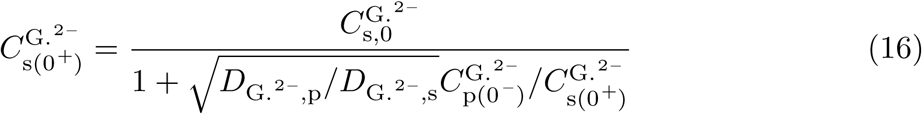

where the ratio 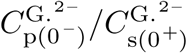 can be obtained from Eq 13c. The expressions in Eq 15 are also approximately valid for short times in a bounded system for times on the order of 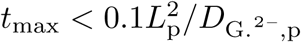 where *L*_p_ is the length of the polyelectrolyte solution. With *L*_p_ = 1 cm and 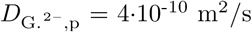, this gives *t*_max_ < 7 h, which should be considered as an order of magnitude estimate above which the effect of the boundaries in the system starts to influence the concentration profile significantly. However, the expressions in Eq 15 give an estimate of the time scales needed for the Gd(DTPA)^2–^ ions to diffuse in the system. The time it takes for the polyelectrolyte solution to reach a certain average value would then be expected to scale as 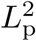.

### Approximate analytic expressions vs finite element simulations

To illustrate under which conditions the approximate relations above are valid, we compare them with FEM solutions for the exact (within the model) equations. Fig 5 shows the concentration profiles using the analytical expressions in Eqs 15 and S18 and how it compares to the simulated values at different times for a low FCD = −50 mM (Fig 5A), for the highest FCD used in this study, i.e. −108 mM (see Fig 5B), and for a higher FCD = −150 mM (see Fig 5C). It can be seen that the approximate analytic expressions reproduce the concentration profiles for values of FCD of −50 mM and −108 mM, respectively, but less so for FCD = −150 mM on account of the linearization of Eq S2. We note that for short times (less than 2 h), it is non-trivial to include the effect of the membrane on the distribution of ions in the different compartments (see the SI for further information). The concentration profile in the polyelectrolyte solution starts to deviate considerably from the analytical expressions when the Gd(DTPA)^2–^ ions have reached the wall of the container containing the polyelectrolyte solution (not shown).

**Fig 5.**
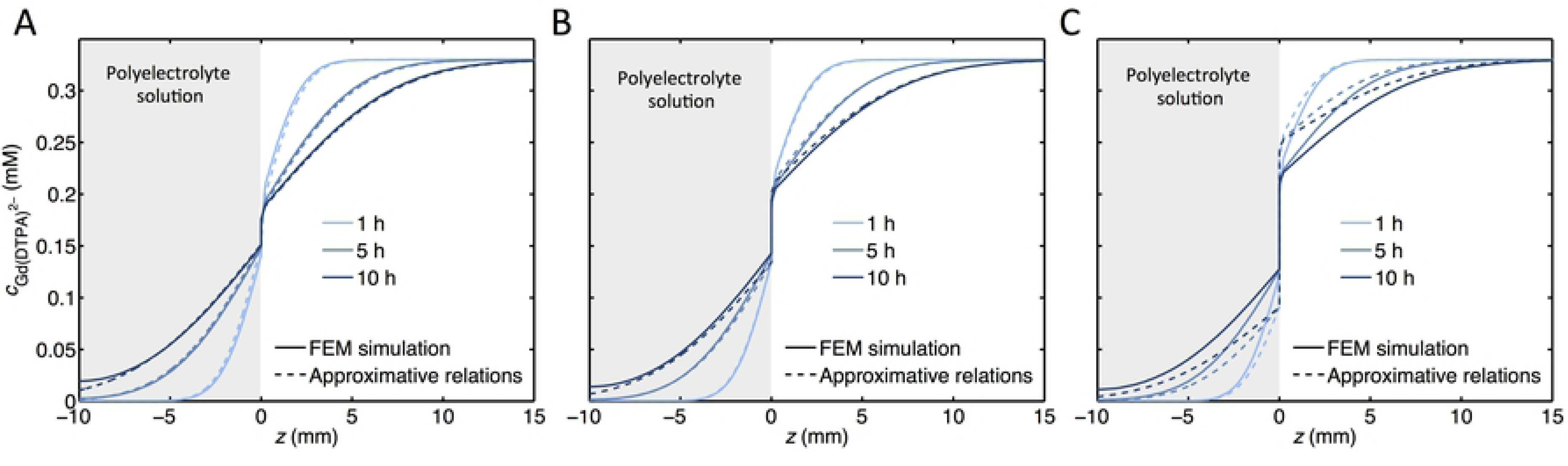
Comparison of concentration profiles from FEM simulations and approximate analytic expressions. The concentration of Gd(DTPA)^2–^ vs *z* from FEM simulations (solid lines) and from Eqs 14 and S18 (dashed lines) for a polyelectrolyte solution with an FCD of **(A)** −50 mM, **(B)** −108 mM and **(C)** −150 mM. Parameters used in the calculations A; 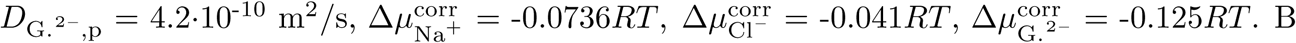. B; values in Table 1. C; 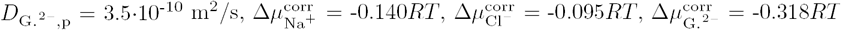.

### Diffusion coefficient of contrast agent in the polyelectrolyte solution

From Eq 15a, it is possible to obtain values of the diffusion coefficient of Gd(DTPA)^2–^ in the polyelectrolyte solution from the experimentally determined concentration profiles. As outlined in the SI, one has to take into consideration the fact that Eq 15a applies to a system with infinite boundaries and that there is convection in the salt solution. When this is accounted for, Eq 15a is modified to Eq S21 in the SI. A fit of Eq S21 to one dataset is given in the SI, and the obtained diffusion coefficients are presented in Table 1. We note that the values follow the expected dependence on the magnitude of FCD. Thus, the values are 91, 87 and 82% of the value in the salt solution for FCD values of −73, −92 and −108 mM, respectively.

## Conclusions

We have combined magnetic resonance imaging results with simulations on a previously developed model system with the aim of studying issues related to early diagnosis and *in vitro* and *in vivo* clinical studies of osteoarthritis. The flow of a charged contrast agent from a salt solution into a polyelectrolyte solution has experimentally been determined with spatial and temporal resolutions of 90 *µ*m and 600 seconds, respectively. The relevant equations describing the flow have been solved using Finite Element Methods. The results show that it is important to take non-ideal conditions arising from molecular interactions into account. Approximate expressions for the flow have been derived and compared with numerical solutions of full equations in order to assess when the approximations are reasonable. Finally, the derived approximate expressions make it possible to determine the diffusion coefficient for the contrast agent in a polyelectrolyte solution from the experimental data.

## Supporting Information

**S1 File. Supporting Information.** Derivation of diffusion equations and relations describing concentration profiles; Donnan equation for the investigated system; Discussion of determination of dialysis time; Additional figures of results from experiments and simulations.

## Acknowledgement

Financial support by the Swedish Research Council (VR) through the Linnaeus grant for the Organizing Molecular Matter (OMM) center of excellence is gratefully acknowledged. PJ was supported by a grant from the Swedish Research Council (number: 621-2014-3907).

